# Biophysical Characterization of the human Nav1.9 sodium channel in trigeminal ganglia and dorsal root ganglia neurons

**DOI:** 10.64898/2026.08.22.746445

**Authors:** Yu Patrick Shi, Tamara Cotta, Ian Orozco, Fanny Chen, Yannick Miron, Richard Kondo, Mark L. Chapman, Douglas S. Krafte, Andre Ghetti, Kevin P. Carlin

**Affiliations:** AnaBios Corporation, San Diego, CA 92101, USA; Niroda Therapeutics, Inc., Short Hills, NJ 07078, USA

**Keywords:** Nav1.9, Trigeminal ganglia, Dorsal root ganglia, Pain

## Abstract

In human dorsal root ganglia (DRG), and trigeminal (TG) neurons, the various voltage-gated sodium channel (Nav) isoforms play critical roles in the firing of action potentials, which drive electrical impulses that encode somatosensations including, itch, and pain. The *SCN11A* gene encodes the tetrodotoxin (TTX)-resistant voltage-gated sodium channel Nav1.9, characterized by unique gating properties. Unlike other isoforms, the Nav1.9 channel activates and inactivates slowly and has a hyperpolarized voltage-dependence of activation and depolarized voltage-dependence of inactivation. This leads to a large window current that has been suggested to function as a regulator of the resting membrane potential of neurons. Mutations in Nav1.9 channels lead to congenital insensitivity to pain (gain-of-function) or familial episodic pain syndrome (loss-of-function) suggesting the channel is a critical mediator of pain. Despite its relevance in pain pathophysiology, most existing data relies on rodent models or heterologous expression systems, leaving the specific pharmacology and biophysical behavior of these channels in human primary neurons largely unknown. In this study, we pharmacologically isolated and characterized native Nav1.9 channel currents in human DRG and TG neurons to compare their biophysical profiles. Our findings reveal significant kinetic and voltage-dependent differences between the two populations. Specifically, Nav1.9 channels in TG neurons exhibit a right-shifted steady-state inactivation curve, a larger window current, and faster activation kinetics compared to those in DRG neurons. In addition, conditions that simulate inflammatory states *in-vivo* greatly potentiates the Nav1.9 currents consistent with similar observations in rodent models. By detailing these distinct biophysical properties, this research offers crucial insights into Nav1.9 channel function relevant for drug discovery efforts aimed at developing analgesics for both acute and chronic pain.

## Introduction

Voltage-gated sodium channels (Nav) are critical in the rapid generation and propagation of action potentials (AP) in neurons that transmit nociceptive information—the sensory signals underlying pain evoked by thermal, mechanical, and noxious chemical stimuli^1^. Of the nine mammalian isoforms of Nav channels (Nav 1.1-1.9), three are highly expressed in dorsal root ganglion (DRG) and trigeminal ganglion (TG) neurons: the tetrodotoxin (TTX)-sensitive Nav1.7 and the TTX-resistant Nav1.8 and Nav1.9^2–11^. As critical players in nociceptive signaling, Nav channels have been pursued as targets for non-opioid analgesics for decades, though with little clinical success. A breakthrough came in early 2025 with the FDA approval of JOURNAVX (VX-548, suzetrigine), a potent, selective Nav1.8 inhibitor from Vertex Pharmaceuticals—which reinvigorated the non-opioid analgesic field and spurred efforts to develop improved compounds against this and related targets^12,13^. Despite a relatively incomplete understanding of the channel, Nav1.9 remains of strong interest in the pain field, owing to its predominant expression in small-diameter DRG and TG neurons and its proposed role in regulating nociceptor excitability and pain signaling^2,10,14–17^. Gain-of-function mutations in the Nav1.9 channel are associated with congenital indifference to pain (CIP)^18,19^, familial episodic pain (FEP)^20,21^, and painful peripheral neuropathy (PPN)^22^, suggesting this channel is an important regulator of pathologic pain and is a promising analgesic target. Structurally, Nav1.9 resembles a typical voltage-gated sodium channel. The gene, *SCN11A*, encodes the α-subunit of Nav1.9, which consists of four homologous domains (DI-DIV) with each domain consisting of six transmembrane helices (S1-S6)^23^. S1-S4 is the voltage sensing domain (VSD), and the S5-S6 is the pore forming domain (PD). Serine at DI-SS2 underlies the TTX-R phenotype, along with the conserved PKC phosphorylation site at L3, and the conserved IFM motif involved in fast inactivation^23^. Despite these structural similarities, Nav1.9 channels demonstrate unique gating properties where activation occurs at more hyperpolarized potentials, and both activation and inactivation rates are significantly slower than other Nav isoforms thus creating a larger window current than that of Nav1.7 and Nav1.8^19,22,24–26^. The slow gating kinetics of Nav1.9 suggest that the channel contributes little to the action-potential upstroke, functioning instead as a modulator of the resting membrane potential (RMP): by generating a sustained, subthreshold depolarizing current, it may sensitize nociceptors and lower the current threshold required to trigger an action potential^27,28^. In addition, the human Nav1.9 channel amino acid sequence is only 75% identical to rodent channel, making Nav1.9 channels the least conserved amongst all the Nav channel orthologs^29^. Indeed, previous biophysical characterization of Nav1.9 in native human DRG neurons revealed that this channel exhibit lower threshold for activation, higher threshold for inactivation, slower deactivation, and faster recovery from slow inactivation compared to mouse DRG neurons^30^.

Until recently, efforts to characterize Nav1.9 were hampered by the channel’s notorious resistance to functional expression in heterologous systems—a barrier that is only now being overcome. Stable cell lines and reliable transient transfection in HEK293 and ND7/23 cells have since been established, enabling both high-throughput screening and mechanistic study^19,25,26,31^.

To enhance and stabilize Nav1.9 currents for these pharmacological studies, recordings are typically supplemented with GTPγS, a non-hydrolyzable GTP analog that persistently activates G-proteins^32,33^. Even so, an important gap remains: Nav1.9 has been studied almost exclusively in rodent neurons and heterologous expression systems, leaving its properties and role in native human cells poorly defined. Only recently did Zhang et al. (2025) report a direct biophysical comparison of human and rat Nav1.9^30^. A further limitation has been the absence of a selective Nav1.9 inhibitor or agonist to serve as a pharmacological tool for probing the channel’s contribution to neuronal excitability. This is beginning to change: Potet et al recently disclosed a relatively selective Nav1.9 compound, prompting several pharmaceutical companies to synthesize their own versions for use as tool compounds^34^.

In the present study, we further characterize native Nav1.9 channels expressed in human DRG neurons and provide the first recordings and biophysical characterization of native Nav1.9 channels in human TG neurons. Our comparative analysis reveals several key biophysical distinctions suggesting tissue-specificity in Nav1.9 channels: native Nav1.9 channel in TG neurons exhibits a right-shifted voltage-dependence of inactivation despite an unaltered activation profile, leading to larger window current, faster activation kinetics, and slower recovery from slow inactivation kinetics compared to DRG neurons. Furthermore, in our study application of a cocktail of inflammatory mediators (inflammatory soup (IS)) significantly potentiated the Nav1.9 current in DRG neurons, contrary to what has been reported previously^30^. IS shifted the voltage-dependence of activation toward more hyperpolarized potentials while having only a minor effect on inactivation. Collectively, these findings suggest that Nav1.9 channels may play a more prominent role in priming resting membrane potentials and increasing excitability in TG neurons compared to DRG neurons. Under inflammatory conditions, modulated Nav1.9 channels are likely to drive both DRG and TG neurons toward a hyperexcitable state.

## Material and Methods

### Human DRG and TG Tissue Procurement

The procurement network of AnaBios Corporation includes only US-based Organ Procurement Organizations and Hospitals. Policies for donor screening and consent are those established by the United Network for Organ Sharing (UNOS). Organizations supplying human tissues to AnaBios follow the standards and procedures established by the US Centers for Disease Control (CDC) and are inspected by the Department of Health and Human Services (DHHS). The distribution of donor medical information is in compliance with HIPAA regulations to protect donors’ privacy. All transfers of donor tissue to AnaBios are fully traceable and periodically reviewed by US Federal authorities. Donor TG and DRGs, from ages: 18 – 60 years old, were recovered using AnaBios’ proprietary surgical techniques and were shipped to AnaBios. The DRGs were then further dissected in a cold proprietary neuroplegic solution to remove all connective tissue and fat. The ganglia were enzymatically digested, and the isolated neurons were cultured in DMEM F-12 (Gemini Bio-Products, Cat. No. 900–955, Lot No. M96R00J) supplemented with glutamine 2 mM, horse serum 10% (Invitrogen, Cat. No. 16050–130), human nerve growth factor (hNGF) (25 ng/ml) (Cell Signaling Technology, Cat. No. 5221LF), GDNF (25 ng/ml) (ProSpec Protein Specialist, Cat. No. CYT-305) and penicillin/streptomycin (ThermoFischer Scientific, Cat. No. 15140–122).

To minimize neurite outgrowth and achieve a well-clamped system for electrophysiology recordings, dissociated DRG and TG neurons were kept in hibernate solution (Fisher Scientific, Cat. No. NC0442869), supplemented with 1% pen/strep, as suspension culture and stored at 4°C prior to plating. On the day of the experiment, the suspension DRG and TG neurons were spun down and resuspended in culture media without growth factors, plated onto 12 mm poly-D-lysine coated glass coverslips and incubated at 28°C supplemented with 5% CO_2_. Both DRG and TG neurons were recorded 1-2 days after plating.

### Chemicals

Tetrodotoxin (TTX, Cat #1069) citrate was purchased from Tocris Bioscience (Minneapolis, MN) and prepared to a 5 mM stock concentration in milli-Q water and diluted 10,000x to a working concentration of 0.5 µM. Suzetrigine (VX-548, Cat #HY-148800) was purchased from MedChemExpress (Monmouth Junction, NJ) and prepared to aliquots of 300 µM stock concentration in DMSO and was diluted 1,000x to a working concentration of 300 nM with external recording solutions. Senktide (Cat #1068) and TC-N 1752 (Cat #4435) were purchased from Tocris Bioscience (Minneapolis, MN). ICA604025 was generously provided by Niroda Therapeutics Inc. Compounds were prepared at 10 mM stock concentration with DMSO, and diluted 1,000x to a working concentration at 10 µM. Inflammatory soup (IS) comprised a cocktail of the following: 1 μM prostaglandin E2 (Tocris Cat #2296), 10 μM of histamine (Sigma-Aldrich Cat #H7250), 1 μM of serotonin (Sigma-Aldrich Cat #9523), and 1 μM of bradykinin (Sigma-Aldrich Cat #B3259). GTP-gamma-S (GTPγS) was purchased from Abcam (Cat #ab146662, discontinued). A 100 mM stock concentration of GTPγS was prepared in milli-Q water and diluted to 0.5 mM in the internal recording solution, unless stated otherwise.

### Electrophysiology

All whole-cell patch clamp recordings were performed at room temperature (20-22°C) with Multiclamp 700A/B (Molecular Devices, Sunnyvale). Signals were filtered at 3 kHz with a low-pass filter and sampled at 10 kHz. Patch pipettes were pulled from borosilicate glass using a P-1000 Sutter puller (Sutter Instruments). Pipette had a resistance of 1-3 MΩ when filled with internal recording solution containing (in mM): 70 Cs-Methansulfonate, 20 CsF, 7 NaCl, 1 MgCl_2_, 0.5 CaCl_2_, 30 TEA-Cl, 10 EGTA, 10 HEPES, 3 Na-ATP, and 0.5 GTPγS, pH to 7.2 with CsOH.

External recording solution contains (in mM): 120 NaCl, 3 KCl, 1 MgCl_2_, 2 CaCl_2_, 0.1 NiCl_2_, 0.1 CdCl_2_, 2 CsCl, 25 Glucose, 10 HEPES, 20 TEA-Cl, 4 4-AP, 0.0005 TTX, and 0.0003 VX-548, pH to 7.4 with NaOH.

To generate the activation conductance curve, cells were held at -80 mV and a prepulse to -100 mV was delivered for 1 s before activating the channels from -100 mV to +30 mV for 250 ms in 10 mV increments. Peak inward currents were measured to generate the current-voltage relationship (I-V), and the conductance-voltage relationship (G-V) was calculated based on the following equation, *G* = *I*/(V_m_ – E_Na_), where *G* is the conductance, *I* is the peak current, V_m_ is the membrane potential, and E_Na_ is the sodium reversal potential. To assess the steady-state voltage-dependence of fast inactivation, the cells were initially held at -80 mV, then stepped from -100 mV to +30 mV for 500 ms, at 10 mV increments, before depolarizing to -20 mV for 100 ms to elicit a tail current to assess the proportion of inactivated channels. Peak tail currents were measured after the -20 mV test pulse to generate the inactivation G-V relationship. Both activation and inactivation G-V relationships were normalized and fit to a Boltzmann equation: *G*=*G_max_*/1+*exp*[(V_1/2_-V_m_)/*k*], where V_1/2_ is the half maximum of activation or inactivation, and *k* is the slope.

The activation kinetics (*tau)* at varying activation voltages were determined using single exponential fits to the rising phase of the current. For rate of inactivation, a single exponential fit was used for the decaying phase of the current, following activation at varying depolarization voltages. To assess the rate of deactivation, cells were initially held at -80 mV follow by a prepulse to -100 mV for 1 s, then depolarized to -20 mV to open the channels for 40 ms, before closing the channels with a steps from -100 mV to -10 mV, in 10 mV increments. The decaying currents were fit with a single exponentials to acquire the *tau* of deactivation. To record the recovery from slow inactivation, the cells were initially held at -80 mV before depolarizing to -20 mV for 3 minutes to allow the channel to enter the slow inactivated state. The cell was then hyperpolarized to a given potential (e.g. -100 mV, -80 mV or -60 mV) to recover the channel, followed by a test pulse to -20 mV for 100 ms, and was delivered at 0.2 Hz. The peak inward current at the -20 mV test pulse was then plotted against time allowed for channel recovery and fitted to a single exponential.

### Immunofluorescence

Primary dorsal root ganglion (DRG) neurons were seeded in 24-well plates and fixed with 4% paraformaldehyde (PFA) in phosphate-buffered saline (PBS), then stored in PBS at 4°C. Samples were blocked for 1 hour at room temperature in a blocking buffer containing 5% horse serum and 0.1% Triton X-100 in PBS.

Following blocking, cells were incubated overnight at 4°C with a mouse monoclonal anti-β-Tubulin III (neuronal) antibody (Cat. No. T8578, clone 2G10, Sigma-Aldrich), diluted 1:2000 in 5% horse serum in PBS. The following day, after three washes with PBS, samples were incubated for 1 hour at room temperature with a Rhodamine Red-X-conjugated donkey anti-mouse secondary antibody (1:250; Jackson ImmunoResearch).

Nuclei were counterstained using NucBlue Fixed Cell ReadyProbes Reagent (Cat. No. R37606, Thermo Fisher Scientific). Fluorescence images were acquired using an Olympus IX83 inverted fluorescence microscope equipped with an X-Cite XYLIS LED illumination system and a filter set containing an AT560/40x single-band excitation filter and an AT635/60m single-band emission filter, using a 10× objective. Images were analyzed using MetaMorph and ImageJ software.

## Results

### Nav1.9 voltage-dependence

To isolate Nav1.9 current, neurons were initially bathed in external solution containing a combination of 500 nM of TTX and suzetrigine (VX-548) at 300 nM, while being held at -80 mV. 120 mM of external NaCl was used to increase the driving force for sodium currents. Next, we supplemented the internal recording solution with 500 µM of GTPγS, which has been shown to potentiate Nav1.9 currents by activating G-protein signaling pathways^25,32,33^. Activation of Nav1.9 currents was induced by first applying a 1-s hyperpolarization step to -100 mV to allow the channels to reach resting state from a holding potential of -80 mV, followed by a series of 250 ms depolarization steps from -100 mV to +20 mV. The peak currents were then plotted against voltage to generate the activation current-voltage relationship. Conductance-voltage (G-V) relationship was calculated based on *G* = I*γ*N*(V_m_-E_Na_); where *G* is the conductance, I is the current, γ is the single channel conductance, N is the number of channels, and V_m_-E_Na_ is the driving force. To analyze the inactivation G-V relationship, neurons were initially held at -80 mV, and a series of 500 ms depolarizing steps from -100 mV to +20 mV was performed to inactivate the channels. This was then followed by a test pulse to -20 mV for 40 ms to generate a tail current. Consistent with previous reports, GTPγS caused a left-shift in the V_1/2_ of activation from -18.9 ± 2.5 mV (without) to -29.9 ± 1.1 mV (with), and the V_1/2_ of inactivation was left-shifted from -24.8 ± 2.7 mV to -30.4 ± 1.1 mV (Figure S1), which is much larger than that observed in heterologous expression system (Figure S1)^25^. Additionally, we found that the currents took more than 5 minutes to potentiate and stabilize after break-in. Unless indicated, all experiments were performed in the presence of GTPγS.

Interestingly, we found that voltage-clamp-based recordings of Nav1.9 currents were especially susceptible to space-clamp issues after the neurons were a few days in culture. These un-clamped currents likely arose from the growing neurites suggesting the possibility of high expression level of Nav1.9 along the developing neurites. Primary DRG and TG neurons, once isolated, are covered with satellite glial cells which typically require 3-5 days to detach and expose the neuronal membrane that allows for patch-clamp experiments. During this time, the neurons exhibit rapid neurite outgrowth when cultured in standard culturing media supplemented with GEM21 and growth factors including NGF, and GDNF. Immunofluorescence, using an anti-beta-tubulin III antibody (TUJ1), showed significant neurite outgrowth within 24 hours of dissociation, isolation, and plating on poly-D-lysine coated coverslip. Extensive outgrowth was observed after 72 hours (*days-in-vitro* DIV3) and 120 hours (DIV5) (Figure 1A, *top*). Due to this extensive neurite network, establishing a well-clamped whole-cell system was extremely challenging. The red trace in Figure 1B shows a poorly clamped recording of a neuron cultured in standard media at less than 24 hours. To reduce neurite outgrowth and ensure adequate space-clamp, isolated neurons were immediately transferred to Hibernate solution and kept in suspension at 4°C until the day of the experiment. On the day of recording, the neurons were plated onto Poly-D-lysine coated coverslips in the absence of GEM21 and growth factors and recordings were performed within 12 hours. Immunofluorescence on primary DRG neurons using this culturing method at under 24 hours, DIV3, and DIV5 showed significant reduction in neurite outgrowth (Figure 1A, *bottom*) and a well-clamped Nav1.9 current can be reliably recorded (Figure 1B, *bottom*). Figure 1 shows representative activation and inactivation current traces recorded in DRG (Figure 1C), and TG (Figure 1D) neurons. In general, these large inward currents were slow to activate and showed relatively large persistent current during the depolarizing voltage steps, consistent with that of a Nav1.9 current. (Figure 1E) Fitting the Nav1.9 channel activation and inactivation G-V relationship with the Boltzmann equation in DRG neurons yields a V_1/2_ of -29.9 ± 1.1 mV, and *k* of 7.2 ± 0.9 (*n* = 21), and a V_1/2_ of -30.4 ± 1.1 mV, and *k* of -8.4 ± 1.1 (*n* = 21), respectively (Figure 1C). In TG neurons, the Nav1.9 channel activation V_1/2_ was -30.8 ± 1.6 mV, and *k* of 7.2 ± 1.4 (*n* = 9), and inactivation -22.0 ± 1.0 mV, and *k* of -5.7 ± 0.8 (*n* = 9). These data show that DRG and TG neurons have comparable V_1/2_ of activation and slope, however inactivation V_1/2_ in TG neuron is significantly right-shifted (Figure 1E, inset. *p* = 0.015). This results in a considerable increase in the window current in TG neurons (Figure 1E). This suggests a tissue-specific difference in Nav1.9 current may exist with Nav1.9 channels in TG neurons having greater channel availability around resting membrane potentials and therefore possibly play a bigger role in controlling the membrane potential and excitability compared to DRG neurons.

**Figure 1:**
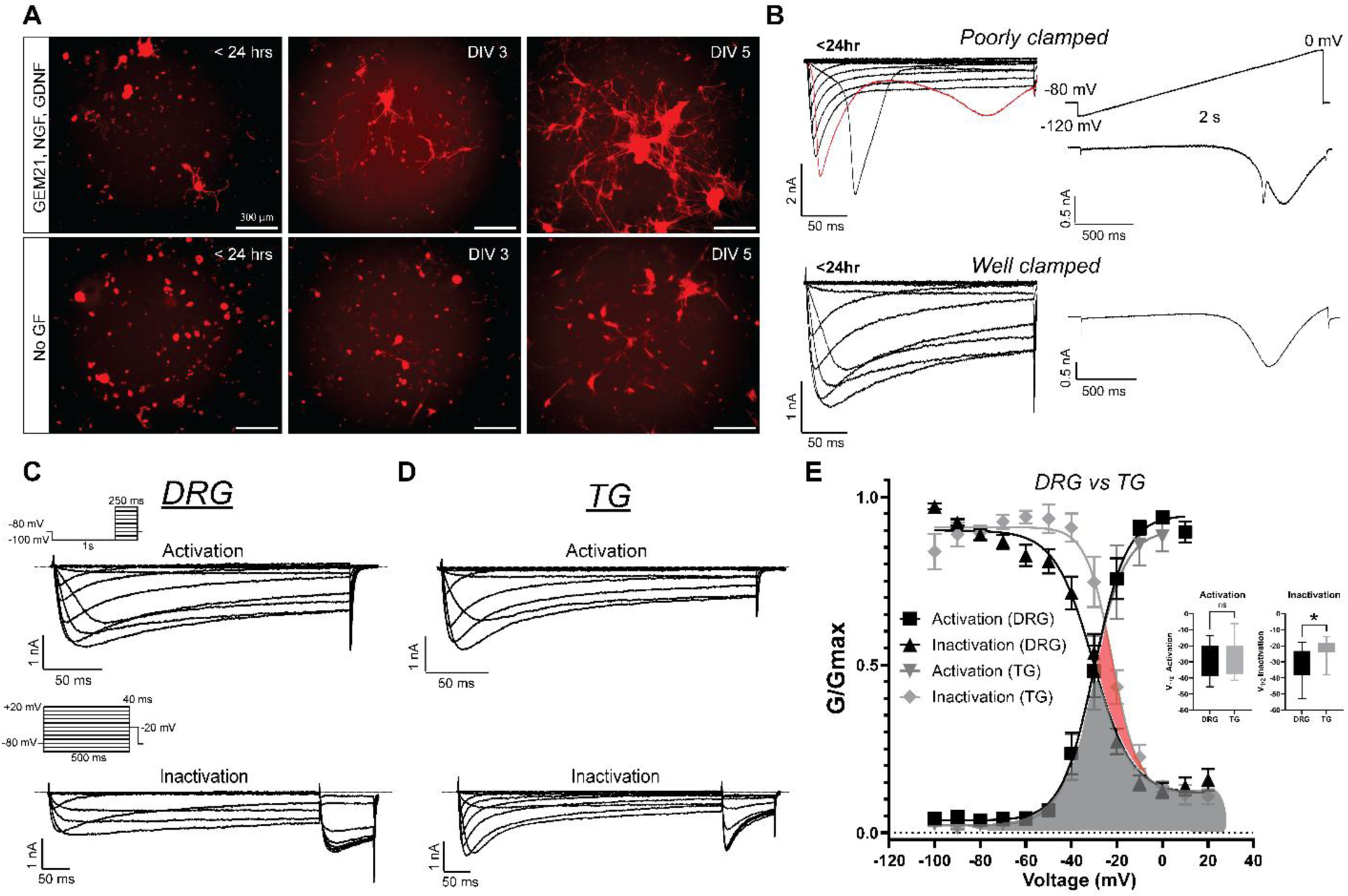
Nav 1.9 channel activation and inactivation properties in native primary human DRG and TG neurons. (**A**) Immunofluorescence of human DRG neurons cultured at 37°C for less than 24 hours, DIV3, and DIV5 in standard DRG culture media (*top*) versus DRG culture media without growth factor (*bottom*). TUJ1 antibody was used to label beta-III tubulin to assess neurite outgrowth. The absence of GEM21, NGF, and GDNF markedly reduced neurite outgrowth leading to significantly better clamped neurons during voltage-clamp experiments. (**B,** *top*) Nav 1.9 currents measured from neurons cultured in standard media at less than 24 hours, showed large distal currents observable in both square voltage step (red trace) and ramp protocol, whereas Nav1.9 currents, measured from neurons cultured without added GF showed well clamped currents (**B,** *bottom*). (**C**) Representative activation and inactivation Nav 1.9 currents recorded in human DRG and (**D**) human TG neurons in extracellular solution consisting of 500 nM TTX, and 300 nM Suzetrigine. (**E**) Activation conductance-voltage (G-V) relationship was elicited by first hyperpolarizing to -100 mV for 1s from a holding potential of -80 mV, then stepping from -100 mV to +20 mV, in 10 mV increments, for 250 ms to activate the channels. In DRG neurons, Boltzmann fit yield an activation V1/2 of -29.9 ± 1.1 mV, and *k* of 7.2 ± 0.9 (*n* = 21). Inactivation G-V relationship was elicited by stepping from -100 mV to +20 mV for 500 ms to inactivate the channel and followed by a test pulse to -20 mV for 40 ms. For DRG neurons Boltzmann fit yields an inactivation V1/2 of -30.4 ± 1.1 mV, and *k* of -8.4 ± 1.1 (*n* = 21). For TG neurons, Boltzmann fit of the activation G-V relationship yield a V1/2 of -30.8 ± 1.6 mV, and *k* of 7.2 ± 1.4 (*n* = 9), and the inactivation G-V relationship yields a V1/2 of -22.0 ± 1.0 mV, and *k* of -5.7 ± 0.8 (*n* = 9). Comparison of the activation and inactivation relationships between DRG and TG reveal that G-V of inactivation in TG is significantly right-shifted (*p* = 0.015) compared to DRG, and shallower in slope. This leads to an increase in window currents, when compared to human DRG (in red area). Taken together these data suggest that the Nav 1.9 current may play a bigger role in human TG and thus inhibition of the channel may be analgesic in neurological pain such as trigeminal neuralgia.

Additionally, the biophysical characteristics of Nav1.9 channels exhibit limited donor to donor variability. Analysis of Nav1.9 activation and inactivation V_1/2_ recorded in DRG neurons across six donors showed that there were no significant differences in the isolated Nav1.9 currents (Figure S2). Some donors exhibited a larger spread in the activation V_1/2_, and a possible explanation is that the currents during the activation protocol are much more prone to being influenced by distal Nav1.9 currents, which can skew the V_1/2_. This further emphasizes the importance of achieving a well-clamped system when recording Nav1.9 currents.

### Nav1.9 Kinetics

Next, we wanted to compare the activation, fast inactivation and deactivation gating kinetics between TG and DRG neurons (Figure 2). Current activation was elicited by depolarizing steps to various potentials from a resting potential of -100 mV then a single exponential fit to the first 20 ms was used to assess the rate of activation (Figure 2A). Nav1.9 currents in TG neurons showed significantly faster activation kinetics between -30 to -10 mV than the currents in DRG neurons (Figure 3A). At -30 mV the average *tau* of activation for DRG and TG neurons were 12.4 ± 1.2 ms (*n* = 5) and 8.2 ± 0.9 ms (*n* = 5), respectively, *p* < 0.02. At -20 mV the DRG and TG neurons had an activation *tau* of 9.2 ± 1.4 ms (*n* = 7) and 5.7 ± 0.70 ms (*n* = 7), respectively, *p* < 0.02. Finally at -10 mV the activation *tau* for DRG and TG neurons were 6.3 ± 0.5 ms (*n* = 7) and 4.3 ± 0.6 ms (*n* = 7), respectively, *p* < 0.02. This suggests that as the membrane depolarizes, Nav1.9 channels in TG neurons activate much faster than in DRG neurons. To assess fast inactivation, the decay of the current was fitted to a single exponential function; TG and DRG neurons show no significant differences in the fast inactivation kinetics (Figure 2B). To assess deactivation of Nav1.9 channels, the neurons were initially held at -80 mV, and after a 1-s hyperpolarization step to -100 mV, a brief 40-ms depolarization pulse to -20 mV was delivered to activate the channels. This was then followed by a series of hyperpolarization steps to deactivate the channel. Single exponential fit to the decay of the tail current in TG and DRG neurons showed no difference in the deactivation kinetics (Figure 2C). Taken together these data show that Nav1.9 channels in TG neurons activate significantly faster than DRG neurons, suggesting the channel passes a bigger current during similar membrane voltage change, and therefore potentially plays a bigger role in the excitability of TG neurons compared to DRG neurons. Furthermore, these faster activation kinetics may allow Nav1.9 currents to contribute more to the action potential waveform in TG neurons, which has been suggested in modeling data^28^.

**Figure 2:**
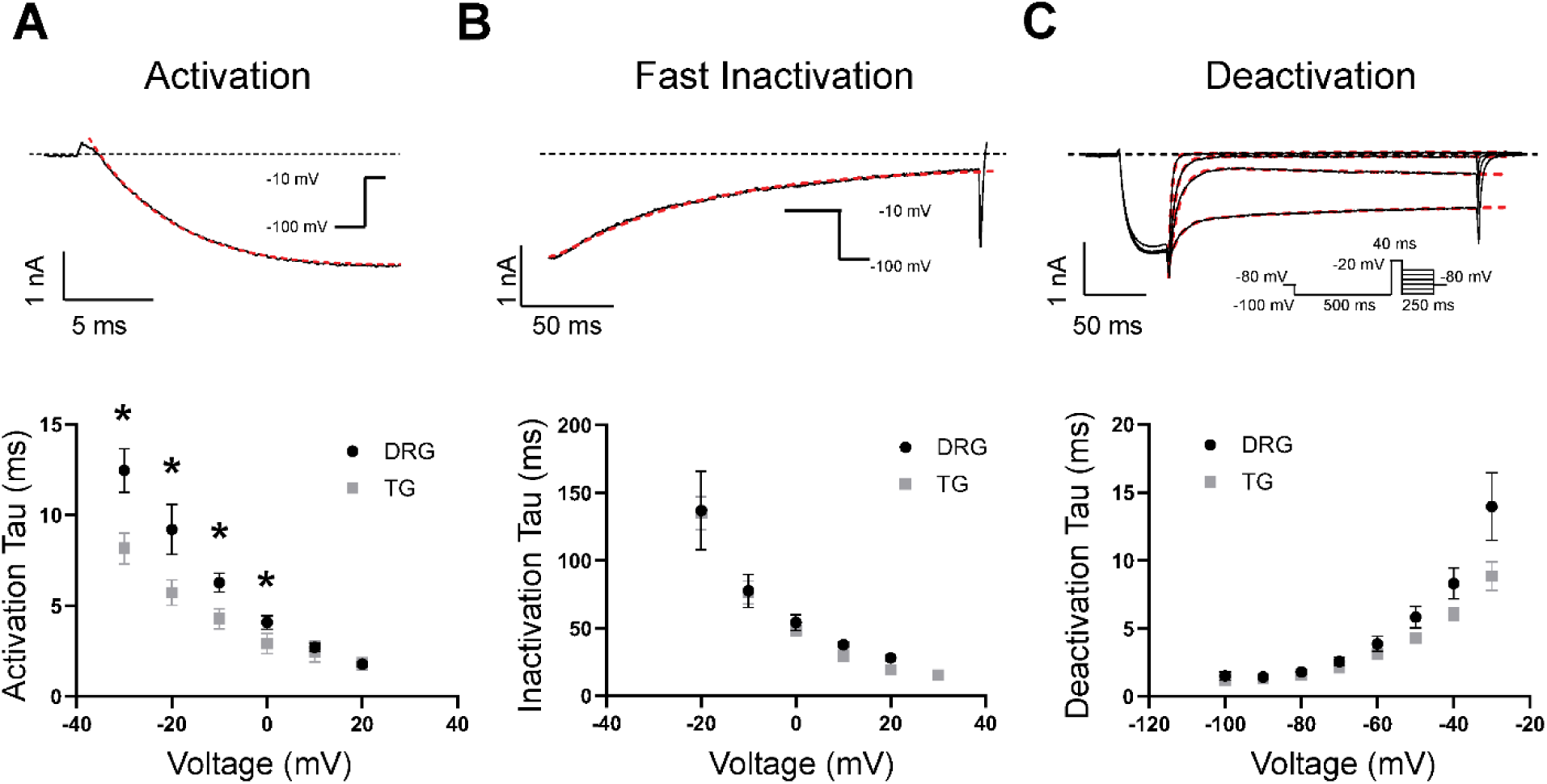
Comparison of Nav1.9 kinetics between human DRG and TG. Nav1.9 channels were activated by initially holding the neurons at -80 mV, then hyperpolarized to -100 mV for 1s before depolarizing to varying potentials. Onset of activation happens rapidly, followed by fast-inactivation. Single exponential fits the onset of current yields activation *tau* (**A,** *n* = 7). Comparison of the activation *tau* indicates that Nav1.9 in TG neurons significantly activates faster at voltages more hyperpolarized than 0 mV. Fit to the decay of the current yields fast inactivation *tau* (**B,** *n* = 7). (**C,** *n* = 7) For deactivation kinetics, the channels were first hyperpolarized to -100 mV for 500 ms, then a brief 40 ms pulse to -20 mV was applied to activate the channel, then followed by hyperpolarization to varying potentials to deactivate the channel. The decay in currents was then fit with a single exponential. There are no significant differences in the fast inactivation and deactivation rate between Nav1.9 channels in DRG and TG neurons. These data suggest that the faster activation rate of Nav1.9 channels in TG neurons would increase the channel availability and potentially sensitize the TG neurons more than DRG.

**Figure 3:**
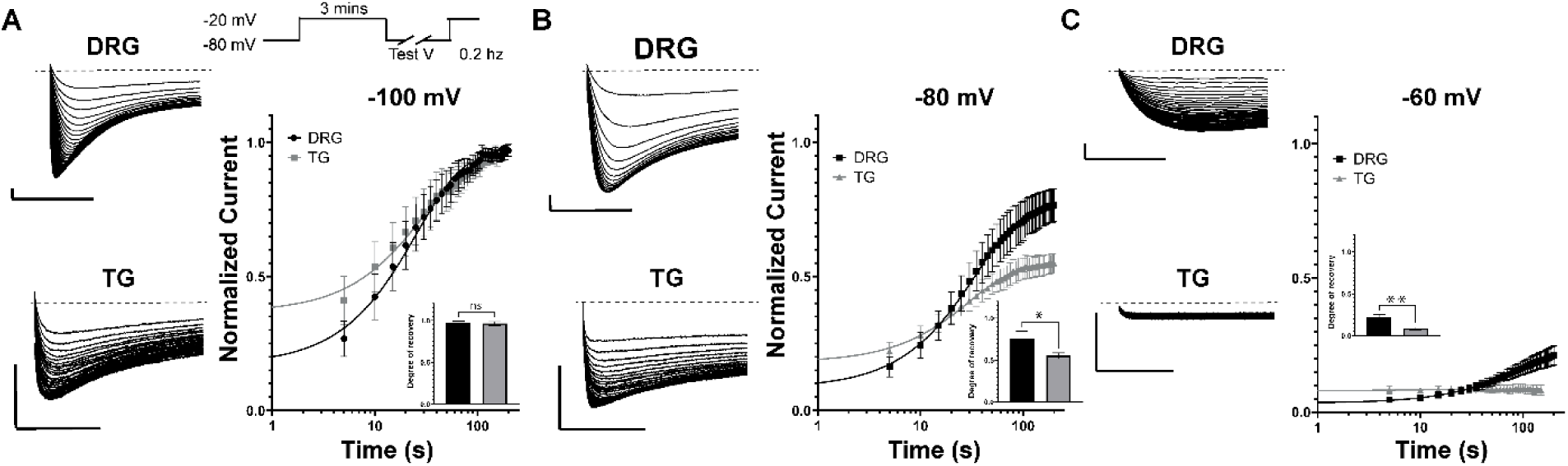
Recovery from slow inactivation between DRG and TG neurons. To assess recovery from slow inactivation, neurons were first depolarized to -20 mV for 3 minutes to drive channels into the slow-inactivated state. Recovery was then monitored at test potentials of -100 mV, -80 mV, or -60 mV, with currents evoked at 0.2 Hz to measure the progressive relief of inactivation. Normalized currents were fitted with a sigmoidal relationship. (**A**) Representative traces from recovery currents of DRG and TG neurons at -100 mV. The recovery from slow inactivation slope constant at -100 mV for DRG neurons was 26.1 ± 2.0 s (*n* = 5) and 30.2 ± 4.7 s for TG neurons (*n* = 5). In both types of neurons, Nav1.9 channel currents nearly recovered to approximately 95% of total. (**B**) Representative traces from recovery currents at -80 mV and the recovery from slow inactivation slope constant for DRG neurons was 35.2 ± 4.1 s (*n* = 5) and 30.7 ± 3.7 s for TG neurons (*n* = 5). The degree of recovery for Nav1.9 channels in TG neurons (55.1 ± 3.7%) was significantly less than DRG neurons (75.6 ± 9.0%) (*inset*). (**C**) Representative traces from recovery currents at -60 mV and the recovery from slow inactivation time constant for DRG neurons was 96.7 ± 32.9 s (*n* = 5). In TG neurons, Nav1.9 channels did not show recovery from slow inactivation within the experimental time frame, suggesting it is significantly slower than that of DRG. In DRG neurons, the channel recovered to 21.9 ± 3.6%. Scale bar represents 50 ms and 1 nA. Data are presented as mean ± SEM.

Lastly, we evaluated the recovery from slow inactivation kinetics between DRG and TG neurons at three different test potentials, -100, -80, and -60 mV (Figure 3). Currents were elicited by depolarizing the neuron to -20 mV for 3 minutes to drive the channels into the slow inactivated state and repolarizing to the test potential at 0.2 Hz before briefly depolarizing to -20 mV to assess proportion of channels that recovered from inactivation. Current-time curves exhibited a sigmoidal relationship and were fitted with a sigmoidal function. At -100 mV both DRG and TG neurons reached nearly full recovery from slow inactivation and no significant differences in the slope of recovery were observed: 26.1 ± 2.0 s (*n* = 5) for DRG neurons, and 30.2 ± 4.7 s (*n* = 5) for TG neurons (Figure 3A). Similarly, there were no statistically significant differences in the slope of recovery at -80 mV for DRG neurons, 35.2 ± 4.1 s (*n* = 5), and TG neurons, 30.7 ± 3.7 (*n* = 5), however the degree of recovery for DRG neurons was close to 76% while only 55% for TG neurons (Figure 3B). Interestingly, at -60 mV, for DRG neurons, 22% of the current recovered and had a recovery time constant of 96.7 ± 32.9 s (*n* = 5), while TG neurons exhibited no recovery from slow inactivation (Figure 3C).

### Nav1.9 Pharmacology

One of the biggest challenges in studying the pharmacology of the Nav1.9 channel is the lack of selective tool compounds. Recently, Potet et al identified and characterized a Nav1.9 selective inhibitor ICA604025^34^. Here, we assessed the pharmacology of Nav1.9 channels, in native human DRG neurons, using both a non-selective state-independent Nav1.9 inhibitor TC-N1752^26,34^, and the selective state-dependent inhibitor ICA604025. Both compounds show strong concentration-dependent inhibition. Figure 4A shows the diary plot of inhibition by TC-N1752 from 10 nM to 10 µM. Currents were elicited by prepulse to -100 mV for 1-s from a holding potential of -80 mV, then a depolarizing step to -20 mV for 40 ms. The Hill fit to the concentration-response relationship yields an IC_50_ of 0.26 ± 0.1 µM (Figure 4B, *n* = 3). Our data is consistent with the previously reported potency of TC-N1752 that was evaluated on hNav1.9 stable cell lines 0.37 µM^34^. Figure 4C, D, showed that ICA604025, tested in the same concentration range, yields an IC_50_ of 0.56 ± 0.9 µM (*n* = 5-6). Our results are comparable to the closed-state IC_50_ reported for recombinant hNav1.9 of 0.76 µM^34^ in HEK293 stable cell line. Taken together, these results help confirm the identity of the slowly activating, persistent sodium currents in these cells and validate that the protocol we used to measure pharmacological effects on Nav1.9 currents in our human DRG neurons is suitable as a pharmacological assay for testing selective Nav1.9 inhibitors.

**Figure 4:**
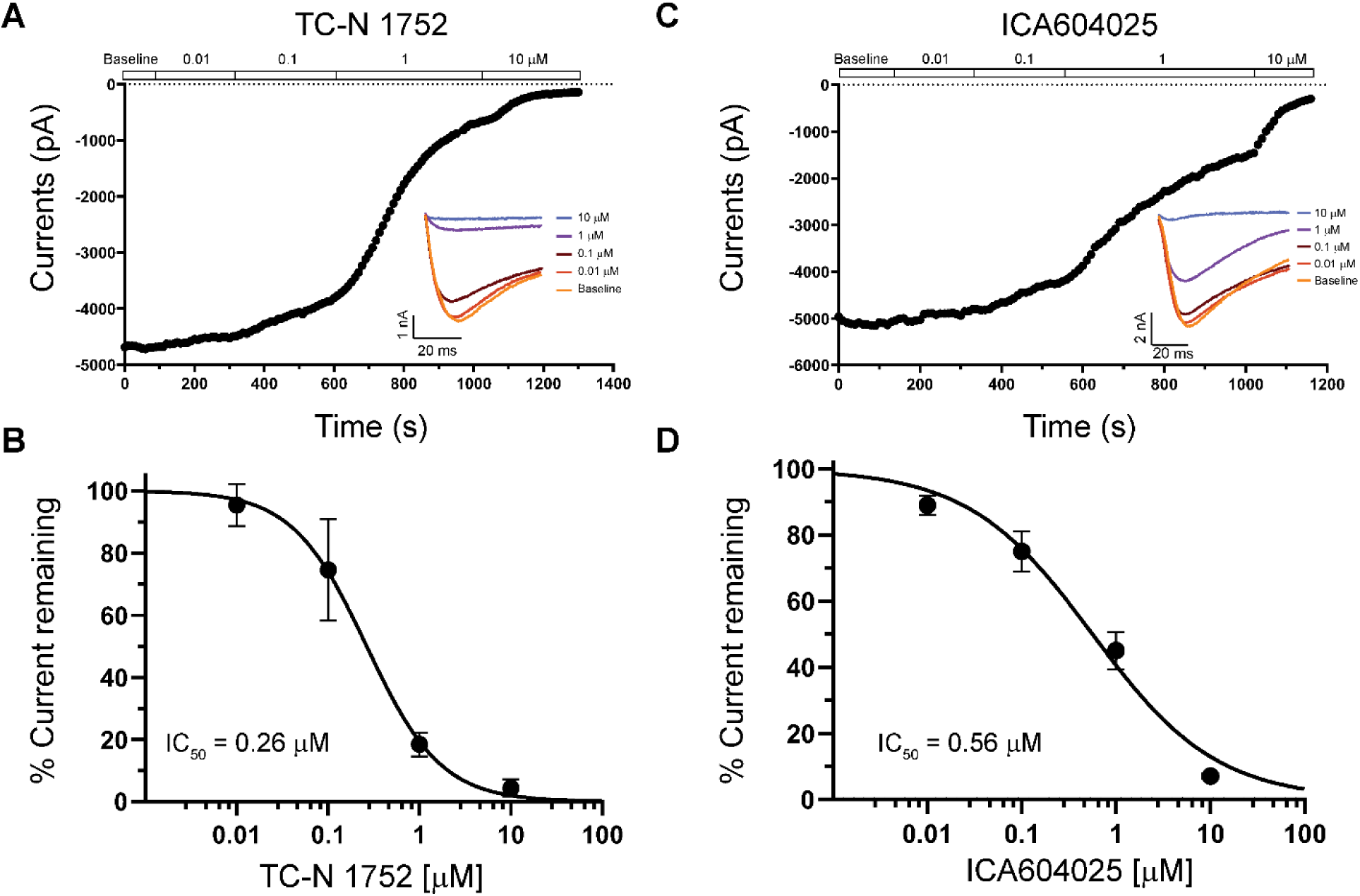
Inhibition of Nav1.9 channels by TC-N1752 and ICA604025 in human DRG neurons . (**A**) TC-N1752 blocked Nav1.9 currents in a concentration-dependent manner from 10 nM to 10 µM reaching full block. Representative traces of Nav1.9 current elicited by first hyperpolarizing to -100 mV for 1 s from a holding potential of -80 mV, then depolarizing to a test potential at 20 mV (*inset*). (**B**) Hill equation fits the dose-response relationship yield an IC50 of 0.26 ± 0.1 µM (*n* = 3), with a slope of -1.1. (**C**) Representative traces of Nav1.9 currents inhibited by increasing concentration of ICA604025, with a strong concentration-dependence. (**D**) Hill fit yields an IC50 of 0.56 ± 0.9 µM (*n* = 5-6), with a slope of -0.7.

TC-N1752 and IC604025 are both small molecule inhibitors that directly interact with Nav1.9 channels to modulate channel gating. A separate mechanism that modulates Nav1.9 gating is through G-protein coupled pathway. Nav1.9 has been shown to be strongly potentiated by G-protein activation. Here, we tested senktide, a potent selective neurokinin-3 (NK3)-receptor agonist, which has been shown to greatly potentiate Nav1.9 currents by causing a hyperpolarizing shift in the voltage-dependence of activation (Figure 5)^35^. Figure 5A shows a representative Nav1.9 current trace recorded from a DRG neurons with varying activation potentials. Application of 10

**Figure 5:**
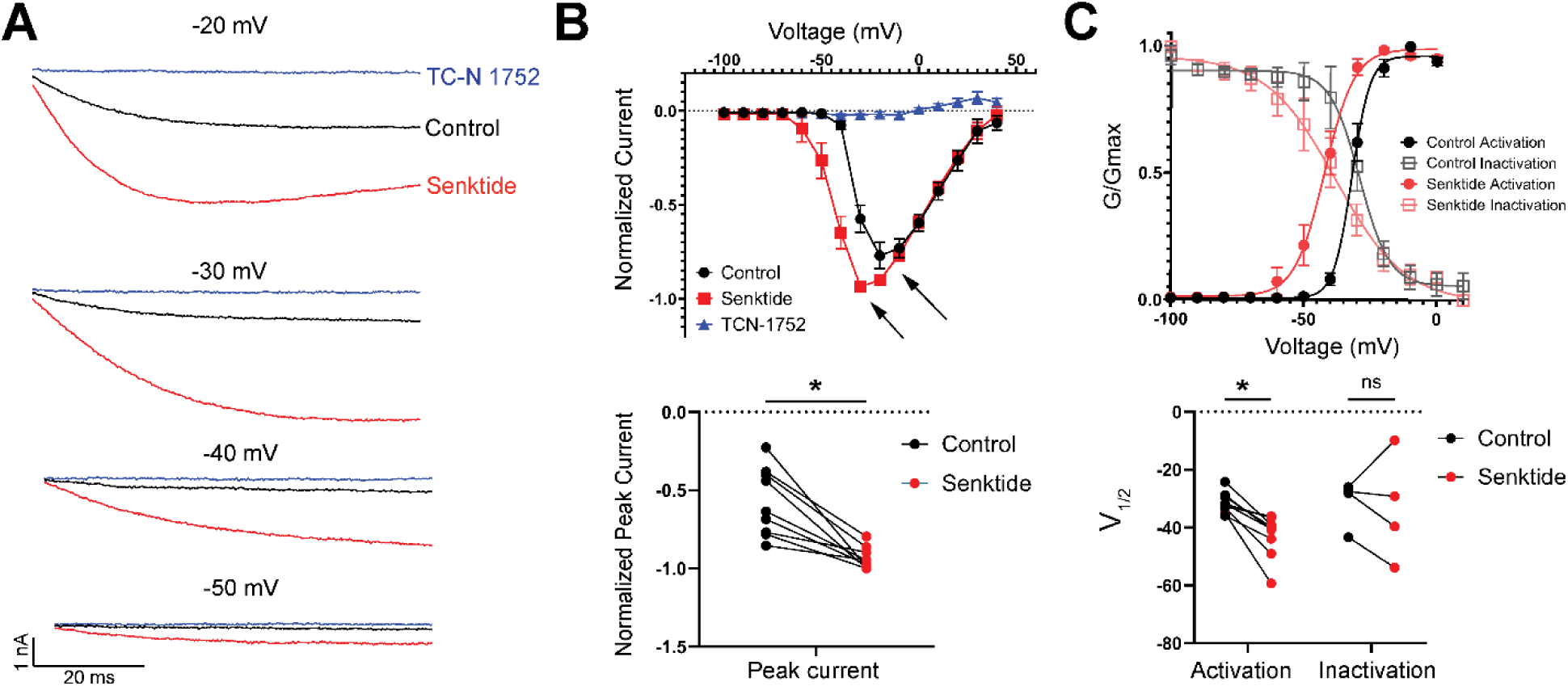
Senktide potentiates Nav1.9 channels in human DRG neurons. (**A**) Representative traces of Nav1.9 currents activated by first hyperpolarizing to -100 mV for 1 s from a holding potential of -80 mV, then depolarizing to test potentials between -20 to -50 mV. 10 µM Senktide potentiated the Nav1.9 currents (red trace), and the non-selective inhibitor TCN-1752 at 10 µM completely inhibited the current (blue trace). (**B**) I-V relationship showed that Senktide significantly increased the peak current (arrow, *p* < 0.01) and induced a hyperpolarization shift in the I-V relationship (*n* = 5), and application of TCN-1752 completely blocked the Nav1.9 currents. (**C**) In the G-V relationship, Senktide caused a significant hyperpolarization shift in the voltage-dependence of activation from V1/2 of -31.4 ± 1.2 mV (*n* = 9) to V1/2 of -42.8 ± 2.4 (*n* = 9, *p* < 0.01) but did not significantly altered the voltage-dependence of inactivation from a V1/2 of -31.3 ± 4.1 (*n* = 4) to V1/2 of -33.2 ± 9.3 (*n* = 4). Data are presented as mean ± SEM.

μM of senktide significantly potentiated the current (red trace), and 10 uM of the non-selective inhibitor TC-N 1752 completely abolished the current (blue trace). From the I-V relationship we observed a ∼8.5 fold increase in currents at -40 mV and a ∼1.2 fold increase in peak current amplitude following application of senktide (Figure 5B, black arrows). In the G-V relationship, the current potentiation is accompanied by a significant hyperpolarizing shift of voltage-dependence of activation by approximately 10 mV from -31.4 ± 1.2 mV (*n* = 9) to -42.8 ± 2.4 mV (*n* = 9; Figure 5C), though voltage-dependence of inactivation and slope factors were statistically unchanged. Our results are consistent with those reported from guinea pig myenteric afferent neurons^35^, though interestingly, the increase in current induced by senktide was observed in the presence of GTPγS, which had already potentiated the Nav1.9 currents. Both senktide and GTPγS are known to activate G-protein signaling, though it is possible that they act through distinct signaling routes that converge to potentiate Nav1.9 (see Discussion). Further investigation is needed to clarify the underlying mechanism^32^.

Evidence primarily from rodent studies indicates that an important pathophysiological mechanism regulating Nav1.9 channel activity, particularly within pain signaling pathways, involves inflammatory mediators^36–39^. In the current study we used an a cocktail of inflammatory mediators: bradykinin, histamine, PGE2, and serotonin^30,36–38,40^ (inflammatory soup (IS)). Application of the IS significantly increased the Nav1.9 currents in human DRG neurons, consistent with previous rodent reports^37,38^, though this is in contrast with recent work in human DRG neurons^30^ (Figure 6A). IS increased the current amplitude by approximately 4-fold when measured at -20 mV and almost doubled the current amplitude and the peak of the I-V curve (Figure 6A). The same effect could be better visualized using a 2-s ramp from -100 mV to +30 mV. Here the IS caused both an increase in peak amplitude and a left-shift in current, resulting in Nav1.9 currents being activated earlier along the ramp (Figure 6B). Consistent with these observations, IS caused a larger shift in voltage-dependence of activation compared to inactivation. Activation voltage-dependence V_1/2_ was left-shifted by 14.2 mV from -12.1 ± 2.9 mV (*n* = 5), to -26.3 ± 2.1 (*n* = 5), however voltage-dependence of inactivation V_1/2_ was left-shifted by only 3.1 mV from -22.4 ± 0.8 mV (*n* = 3), to -25.5 ± 0.4 mV (*n* = 3), though still statistically significant (Figure 6C). Taken together, these data are consistent with what is observed in model systems such as rodent and heterologous expression systems and suggest that potentiation of Nav1.9 currents by inflammatory mediators may play a role in the generation and/or maintenance of pain sensations in pathological states^36–38,40^. Evaluating selective inhibitors against this modulated Nav1.9 current (as in the inflamed state) may provide a more accurate assessment of their analgesic efficacy.

**Figure 6:**
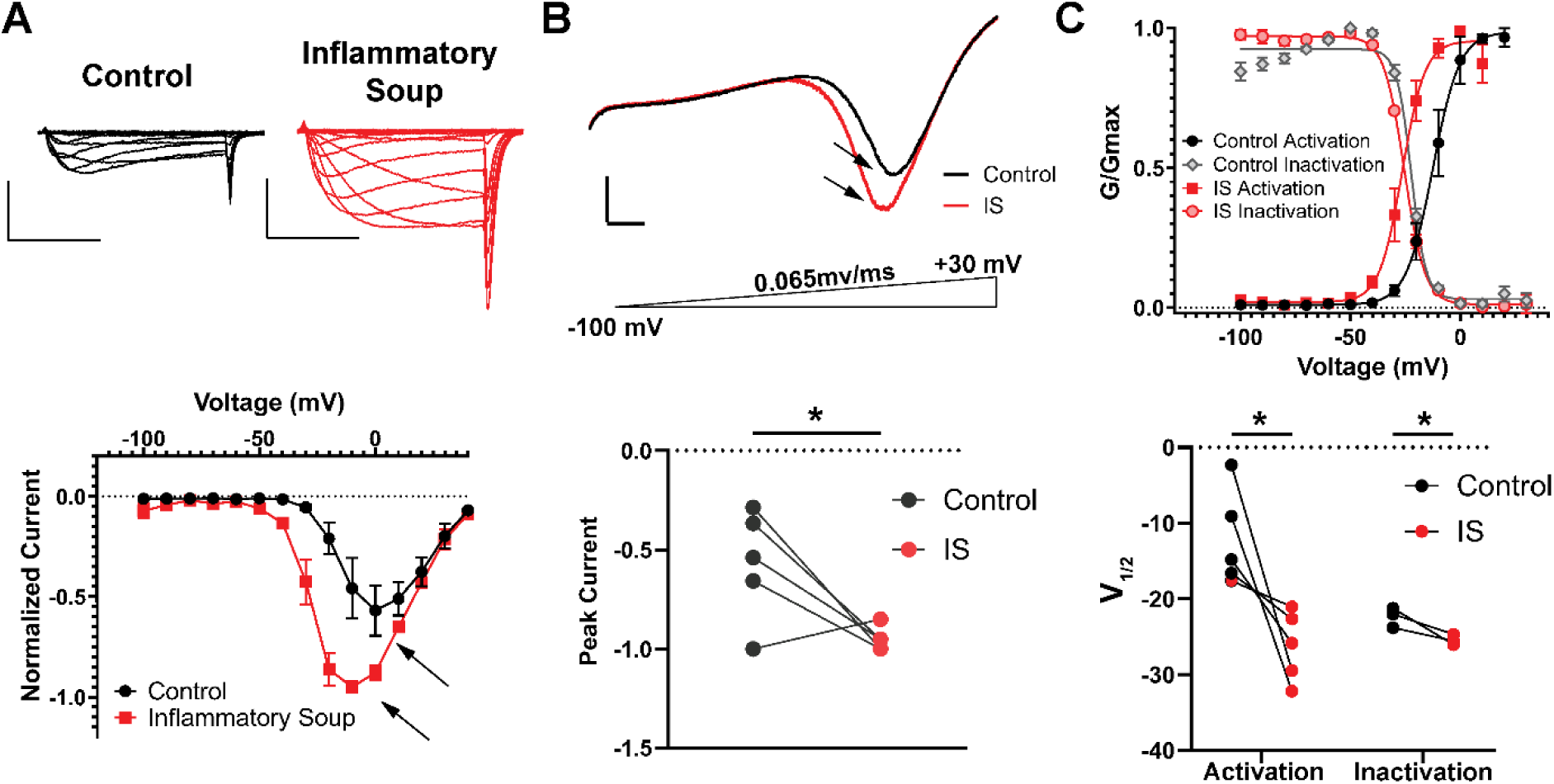
Inflammatory mediators potentiate Nav1.9 currents in human DRG neurons. (**A**) Representative Nav1.9 currents elicited by first hyperpolarization to -100 mV for 1 s from a holding potential of -80 mV, and depolarizing to test potentials between -100 mV to +30 mV for 40 ms. Scale bars represent 20 ms and 2 nA. (**B**) I-V relationship showed that inflammatory soup greatly potentiated the peak current amplitude (arrow) of Nav1.9 currents and induced a hyperpolarization shift in the activation. (**C**) Ramp at 0.065 mv/ms from -100 mV to +30 mV confirmed that inflammatory soup significantly increased the peak current amplitude from 57% to 95% (**D**, *p* < 0.01). Scale bars represent 20 ms and 0.5 nA. (**E**, **F**) G-V relationship showed that the IS significantly left-shifted the voltage-dependence of activation from V1/2 of -12.1 ± 2.9 mV to -26.3 ± 2.1 mV (*p* = 0.005, *n* = 5), and voltage-dependence of inactivation from V1/2 of -22.4 ± 0.8 mV to -25.5 ± 0.4 mV (*p* = 0.02, *n* = 3). Data are presented as mean ± SEM.

## Discussions

In this study, we characterized the native Nav1.9 current in human dorsal root ganglia neurons and for the first time the Nav1.9 current in trigeminal ganglia neurons. Importantly, we identified several differences in biophysical properties of the channels expressed in these two tissues, suggesting tissue specificity in Nav1.9 channels. Before any comparisons could be made, we first needed to establish adequate voltage control of this current in order to make accurate biophysical measurements. However, in both human DRG and TG neurons, Nav1.9 currents proved highly susceptible to space-clamp error: currents arising from electrotonically distant membrane escaped adequate voltage control and contaminated the somatic recording thus obscuring accurate measurement of the current. We postulated that these un-clampable, TTX and VX-548-resistant currents likely arise from Nav1.9 channels trafficked to and expressed along neurites extending from the soma.

Previous reports suggest that Nav1.9 channels contribute to axon elongation in mouse motor neurons, where they act as a trigger for spontaneous Ca^2+^ transients that modulate the rate of axon elongation. Motor neurons from Nav1.9 knockout mice showed both of a loss of spontaneous Ca^2+^ transient, and a decrease in axon elongation^41^. Another report using immunofluorescence showed that contactins in rat DRG directly bind to Nav1.9 channels, and when co-expressed with contactins, greatly enhanced surface expression of Nav1.9^42^. These contactins are expressed along the axons and the highly expressed Nav1.9 channels at these axon locations can possibly produce large distal currents^42^. It is possible that in human DRG and TG neurons, Nav1.9 channels function in a similar manner. It appears as though these channels are rapidly expressed after cell dissociation/plating and are transported along the developing neurites to perhaps act as a modulator for Ca^2+^ transients to drive neurite growth. However, the role and mechanism of Nav1.9 channels in the developing neurites of DRG and TG neurons remain unclear. Nonetheless, to reduce space-clamp issues for the current experiments, we optimized culture conditions that limited neurite outgrowth (such as keeping neurons in suspension, culture without added growth factors, plating cells sparsely, maintaining plated cells at 28°C and recording within 24 hours after plating). These modifications led to well-clamped somatic Nav1.9 currents with minimal contribution from extra-somatic channels. Additionally, we observed that when recording early in culture (<24 hours in-vitro), the majority of DRG and TG neurons expressed robust Nav1.9 currents, irrespective of neuron size, consistent with what has been previously reported^30^. Lastly, we observed very little donor-to-donor variation in the biophysical properties of Nav1.9 channels (Figure 2).

In our electrophysiology recordings, we used methanesulfonate rather than fluoride as the principal internal anion to assess the channel under more physiologically-relevant conditions but with added recording stability over Cs-Cl. Consistent with this rationale, when we used a Cs-F based pipette solution in human DRG neurons, the fluoride produced a 19.5 mV depolarizing shift in the voltage dependence of activation and a 26.7 mV shift in inactivation (Figure S3). This choice came with a trade-off: without fluoride’s seal-stabilizing effect, we could not hyperpolarize to extremely negative potential without compromising seal resistance. Our values using methanesulfonate are consistent with previous reports: the V_1/2_ of activation and inactivation we measured in human DRG neurons are comparable to those reported in rodent DRG neurons and Nav1.9 channels in heterologous expression systems recorded without fluoride^34,43,44^. Under these recording conditions, key biophysical difference between native Nav1.9 channels in human DRG and TG neurons were seen in the voltage-dependence of inactivation. In TG neurons, the voltage dependence of inactivation is shifted approximately 10 mV in the depolarizing direction relative to DRG neurons, with little difference in the voltage-dependence of activation. This effectively widens the window current in TG neurons, which may have significant effects on resting membrane potential and excitability in TG neurons. There are also differences in this biophysical parameter when compared to rat TG neurons. The native Nav1.9 channels in human TG neurons showed markedly depolarized activation and inactivation voltage-dependence relative to rat: the V_1/2_ of activation was −31 mV and -58 mV in rat TG neurons (an approximately 24 mV depolarizing shift) and the V_1/2_ of inactivation was −22 mV and -54 mV in rat TG neurons (an approximately 30 mV depolarizing shift). Also of interest, in human TG neurons the V_1/2_ of activation and the V_1/2_ of inactivation differed significantly from each other, whereas in rat they did not^45,46^.

Nav1.9 channels in TG neurons also appear to activate significantly faster than those in DRG neurons-particularly at hyperpolarized potentials. This suggests that at a given rate of membrane potential change, Nav1.9 channels will pass more current earlier in TG neurons, potentially accelerating the approach to threshold and driving the membrane potential toward more depolarized values. Amongst voltage-gated sodium channels, the relatively slow activation and inactivation kinetics of Nav1.9 channels have led to the proposal that this channel does not directly shape the action-potential waveform, but instead regulates the resting membrane potential to prime nociceptors for firing^4,28,35^. Within this framework, the larger window current and faster activation kinetics we observed in TG neurons would predict a greater Nav1.9 contribution to the resting membrane potential, lowering the threshold for action-potential firing and thereby sensitizing these nociceptors. Moreover, if the modeling data (Kӧster et al) is correct and Nav1.9 current is capable of significantly contributing to the upstroke of the action potential in some sensory neurons, selective inhibition or knockout of Nav1.9 in TG neurons would be expected to reduce excitability and produce analgesia^28^. This would make Nav1.9 a promising drug target for trigeminal neuralgia—a condition so painful it has been termed the “suicide disease”^47^. Consistent with a central role for Nav1.9 in priming TG neurons toward hyperexcitability, Lulz et al. (2015) reported that Nav1.9⁻/⁻ mice show attenuated mechanical and thermal orofacial hypersensitivity following infraorbital nerve constriction, indicating that Nav1.9 is required for the development of orofacial neuropathic pain—though the precise mechanism still remains unclear^27^. Notably, these result contrasts with somatic neuropathic pain models, where Nav1.9 shows no critical involvement, suggesting a trigeminal-specific role that further motivates characterizing the channel in human TG neurons^48,49^.

Recently, a selective Nav1.9 channel inhibitor (ICA604025) was identified and characterized^34^, which paves the way for studying pharmacology in this channel, at least in human and/or nonhuman primate systems where differences in species-dependent pharmacology do not complicate interpretation. Here, our evaluation of the potencies for TC-N1752 (IC_50_ of 0.26 µM) and ICA604025 (IC_50_ of 0.56 µM) against Nav1.9 channels in native human DRG neurons are consistent with previous reports. Previous reports of TC-N1752 potencies were 1.6 µM^26^ and 0.37 µM^34^ and ICA604025 at resting-state potency were 0.76 µM all recorded in stable Nav1.9 cell lines^34^. By refining our experimental and culturing conditions, alongside resting-state pharmacological assays, we reliably generated stable and robust Nav1.9 currents human DRG and TG neurons. These results validate the compound potencies observed in overexpression systems, underscoring the necessity of using native human neuronal platforms to accurately evaluate pharmacological responses under physiological and pathophysiological conditions *in-vivo*. Additionally, we have also evaluated the modulation by a non-selective potentiator senktide, a NK3r agonist on our native human DRG neurons. Senktide both increased the peak amplitude but also induced a hyperpolarization shift in the voltage-dependence of activation similar to that observed in Nav1.9 channels in guinea pig intrinsic primary afferent neurons (IPANS) ^35^. In the same report, it has been suggested that senktide potentiates Nav1.9 current via the PKC pathway and intracellular dialysis of a PKC antagonist, chelerythrine, has been shown to completely abolish senktide-induced potentiation^35^. Interestingly, we see this effect of senktide in the presence of GTPγS in our internal solution, which non-selectively activates G-protein coupled pathways^32,33^. GTPγS potentiation of Nav1.9 currents has been shown to be diminished in the presence of a non-selective protein kinase inhibitor H7 and with ATP removed, suggesting the involvement of protein kinases^32^. One explanation is that GTPγS activates PKA through G_i_ and G_s_ and to a lesser degree via G_q/11_, and the addition of senktide further activates the PKC pathway leading to further current enhancement. However, the exact mechanism of how G-protein pathways potentiate Nav1.9 is still not completely clear.

Lastly, it is well known that inflammatory mediators enhance Nav1.9 currents via the G-protein coupled pathway^25,32,33,36–40^. Here, in native human DRG neurons we find that inflammatory mediators significantly increased peak current accompanied by a hyperpolarization-shift in the voltage-dependence of activation mimicking the effect by senktide and GTPγS (Figure S1), likely through a similar mechanism. Our report is consistent with previously published work from rodent models^37,38,40^. However, this observation is in contrast with a recently published characterization of Nav1.9 in native human DRG neurons, where inflammatory mediators showed a decrease in Nav1.9 currents and no observable shift in the voltage-dependence of activation^30^. A possible explanation to the discrepancy is that our composition of inflammatory soup consists of 10 µM of histamine, which has been shown to enhance Nav1.9 current amplitude, and left-shift the voltage-dependence of activation expressed in ND7/23 cells^50^, a finding consistent with our human data. Differences in the concentrations of the inflammatory mediators used may also play a role in these contrasting results. Nonetheless, the significant increase in Nav1.9 currents and left-shift in voltage-dependence of activation would be expected to increase excitability of both human DRG and TG neurons similar to that described in rodent models^38^. Given the biophysical differences between TG and DRG neurons described, it is possible that in the inflamed state, TG neurons would be even more excitable given the already larger window currents and faster activation kinetics. Our data suggests that Nav1.9 plays an important role in inflammatory-induced hypersensitivity, although further studies in human sensory neurons will be needed to better understand the underlying mechanisms.

## Conclusions

In this study we provide an initial characterization of native Nav1.9 currents in human TG neurons and expanded on the biophysical characterization of this current in human DRG neurons. Our work also provides a novel cell processing and culture method that facilitates the isolation of Nav1.9 currents in human sensory neurons. We found that Nav1.9 channels in TG neurons have a depolarization-shifted voltage-dependence of activation, a larger window current, a faster activation kinetics, and a slower recovery from slow inactivation compared to DRG neurons. These differences suggest that Nav1.9 channels in human TG neurons may play a larger role in neuronal excitability during normal and pathophysiological states such as inflammation. Indeed, our observation that the inflammatory mediators significantly increase the peak current and altered the voltage-dependence of activation of Nav1.9 channels would support this hypothesis. However, the mechanism by which the Nav1.9 channel is being modulated, how much impact Nav1.9 plays in cellular excitability, and the target-ability for developing analgesic are still largely unclear. With the advent of selective Nav1.9 inhibitors, many of these challenging questions will be answered. The kinetic and steady-state voltage-dependence data reported here, obtained from human DRG and TG neurons, may serve as a useful supplement to existing action potential models. By integrating these observations, future computational studies may better evaluate how subtle cellular differences and pharmacological interventions influence action potential morphology and excitability, potentially contributing to the broader effort of designing effective analgesics^28^.

## Acknowledgements

The authors would like to thank the entire tissue procurement team and tissue preparation team at AnaBios Corporations for extracting, dissociating, isolating, and culturing the DRG and TG neurons for patch-clamp and imaging experiments.

## Data availability

Data supporting the findings of this study are available on request from the corresponding author.

## Authorship Contributions

YPS, AG, KC. for conceiving the project. YPS, TC, IO, FC, YM, KC. for methodology, data collection and formal analysis. YPS. for writing - original draft. YPS, TC, IO, FC, YM, RK, MLC, DSK, AG, KC. for writing - review, and editing.

## Conflict of interest

There is no conflict of interest.

## Notes

### Competing Interest Statement

The authors have declared no competing interest.

