## Supplementary Figure for "Biophysical Characterization of the human Nav1.9 sodium channel in trigeminal ganglia and dorsal root ganglia neurons"


**Running title**:

Characterization of Nav1.9 in human TG and DRG neurons


**Address for correspondence:**

Kevin P Carlin
AnaBios Corporation
1155 Island Ave, Suite 200
San Diego, CA, 92101
Phone:




**Keywords**
Nav1.9, Trigeminal ganglia, Dorsal root ganglia, Pain

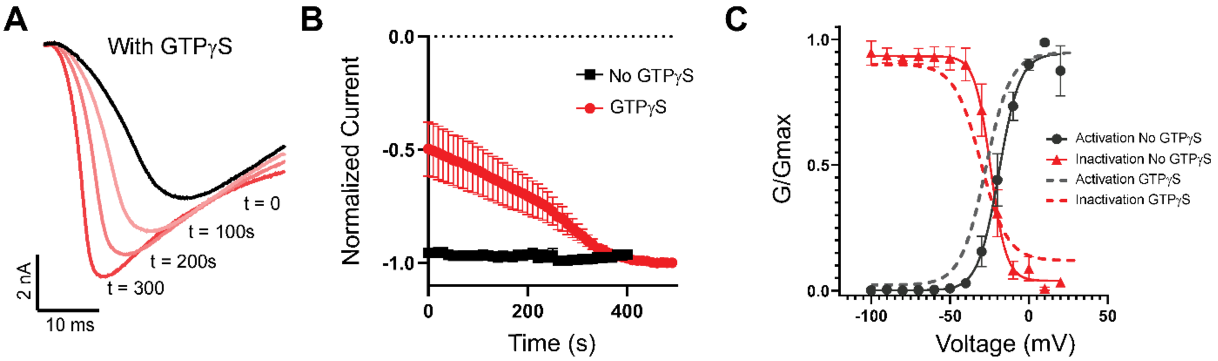


**Figure S1: GTPγS potentiates Nav1.9 currents in human DRG neurons.** (**A**) Representative traces of Nav1.9 currents elicited by depolarizing to -20 mV from -100 mV for 40 ms with a 10 s inter-sweep interval. (**B**) Activation of Nav1.9 currents with internal solution, containing GTPγS, showed gradual potentiation as the neuron is being dialyzed and reached max in ~6 minutes. In contrast, Nav1.9 currents did not show potentiation in the absence of GTPγS in the internal solution. (**C**) GTPγS hyperpolarize-shifted the voltage-dependence of activation from -18.9 ± 2.5 mV (black filled circle, $n = 5$) to -29.9 ± 1.1 mV and hyperpolarize-shifted the voltage-dependence of inactivation from -24.8 ± 2.7 mV (red filled triangle, $n = 5$) to -30.4 ± 1.1 mV. Data was presented as mean ± SEM.

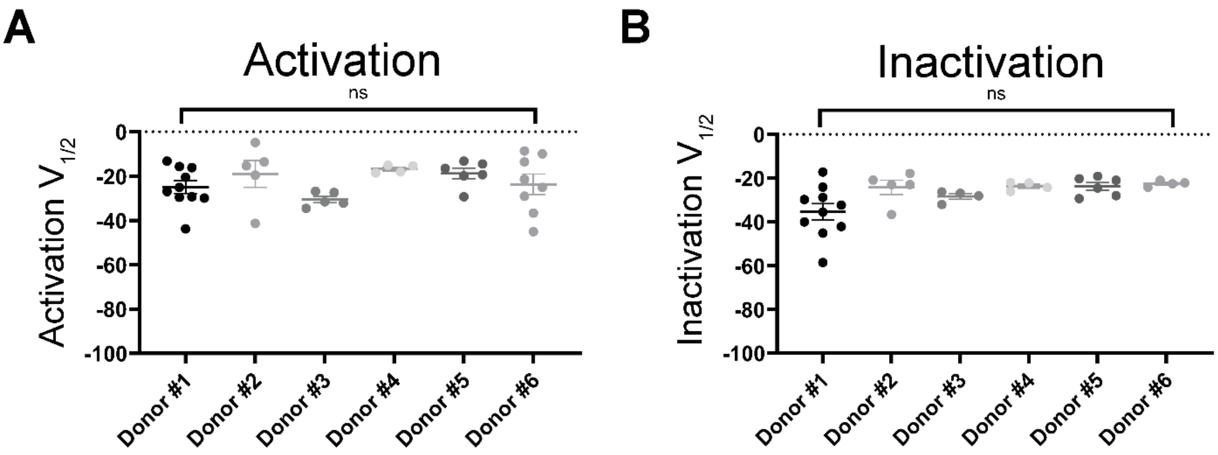


**Figure S2: Nav1.9 currents showed no donor specificity.** Nav1.9 channel activation (**A**), and inactivation (**B**) $V_{1/2}$ measured from individual neurons were plotted from their respective donors. While the activation $V_{1/2}$ values have bigger spread than inactivation $V_{1/2}$, the overall data indicates that there are no significant differences in the biophysical properties of Nav1.9 channels across different donors.

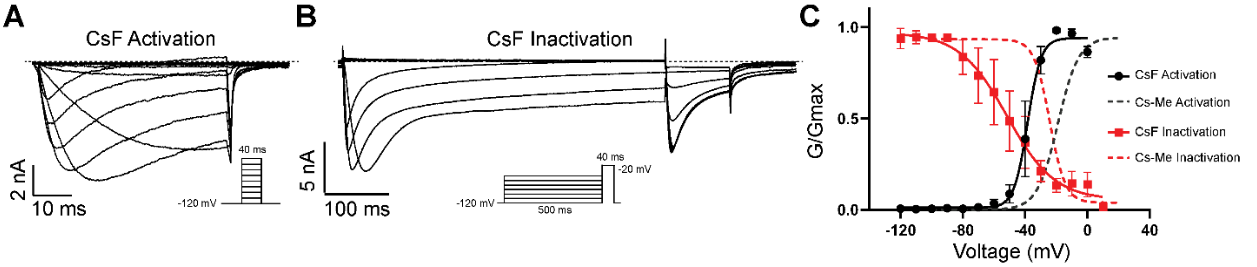


**Figure S3: Cesium Fluoride altered the voltage-dependence of activation and inactivation in Nav1.9 in human DRG neurons.** (**A, B**) Representative traces of Nav1.9 activation and inactivation currents elicited by voltage-clamp protocol shown in the inset. Recordings were done in the absence of GTPγS (**C**) Cesium fluoride-based (CsF) internal solution caused a large hyperpolarization shift in the voltage-dependence of activation and inactivation in comparison to a cesium methanesulfonate-based (Cs-Me) internal solution. Voltage-dependence of activation shifted from a $V_{1/2}$ of -18.9 ± 2.5 mV (black dotted line) to -38.4 ± 0.8 mV (black filled circle, $n$ = 4), and voltage-dependence of inactivation shifted from a $V_{1/2}$ of -24.8 ± 2.7 mV (red dotted line, $n$ = 5) to -51.5 ± 3.5 mV (red filled square, $n$ = 4). Fluoride also caused a substantial shallowing in the inactivation slope from $k$ = -5.0 ± 0.6 in Cs-Me to $k$ = -15.3 ± 4.0. Data were presented as mean ± SEM.
